# Entraining movement-related brain oscillations using rhythmic median nerve stimulation

**DOI:** 10.1101/2020.03.30.016097

**Authors:** Barbara Morera Maiquez, Georgina M. Jackson, Stephen R. Jackson

**Affiliations:** School of Psychology, The University of Nottingham, UK; School of Medicine, The University of Nottingham, UK; Institute of Mental Health, Nottingham, UK

**Keywords:** Median nerve stimulation, Beta entrainment, EEG, Tourette syndrome

## Abstract

Non-invasive brain stimulation techniques delivered to cortical motor areas have been shown previously to: modulate cortical motor excitability; entrain brain oscillations; and influence motor behavior; and have therefore attracted considerable interest as potential therapeutic approaches targeted for the treatment of movement disorders. However, these techniques are most often not suitable for treatment outside of the clinic, or for use with young children. We therefore investigated directly whether rhythmic pulses of median nerve stimulation (MNS) could be used to entrain brain oscillations linked to the suppression of movement. Using electroencephalography techniques together with concurrent MNS we demonstrate that 10 pulses of rhythmic MNS, delivered at 19Hz, is sufficient to entrain Beta-band brain oscillations within the contralateral sensorimotor cortex, whereas 10-pulse trains of arrhythmic MNS does not. This approach has potential in our view to be developed into a non-drug therapeutic device suitable for use outside of the research laboratory or the clinic with brain health conditions associated by excessive movements.

## Introduction

Many brain health conditions, including Parkinson’s disease (PD) and Tourette syndrome (TS), have been linked to the occurrence of unwanted movements and are associated with alteration in the balance of excitatory and inhibitory influences within key brain networks (Ramamoorthi & Lin Y, 2011; Marín, 2012). For instance, TS is a neurological disorder of childhood onset that is characterised by the occurrence of motor and vocal tics, which are involuntary, repetitive, stereotyped movements and vocalisations that occur in bouts, typically many times in a single day (Cohena, Leckman, Bloch, 2013). TS is associated with: dysfunction within cortical-striatal-thalamic-cortical (CSTC) brain circuits that are implicated in the selection of movements (Albin & Mink, 2006); impaired operation of inhibitory (GABA-mediated) signalling (Kalanithi PS, Zheng W, Kataoka Y, DiFiglia M, Grantz H, Saper CB, et al., 2005); and hyper-excitability of sensorimotor regions of the brain, that may contribute to the occurrence of tics (Orth, 2009).

Non-invasive brain stimulation (NIBS) techniques such as transcranial magnetic stimulation [TMS] or transcranial electrical stimulation [tES] have been shown to: alter brain excitability for sustained periods after stimulation [Nitsche, M.A., Paulus, W., 2000]; alter local concentrations of the main excitatory (glutamate) and inhibitory (GABA) neurotransmitters in the brain [Kim, Stephenson, Morris, Jackson, 2014; Stagg, Stephenson, O’Shea, Wylezinska, Kincses, Morris, et al., 2009]; entrain brain oscillations (Pogosyan, Gaynor, Eusebio, Brown, 2009; Thut, Veniero, Romei, Miniussi, Schyns, Gross, 2011); reduce tremor in patients with PD (Brittain, Probert-Smith, Aziz, Brown, 2013); and reduce tics in patients with TS (Kwon, Lim, Lim, Lee, Hyun, Chae, Paik, 2011; Le, Liu, Sun, Hu, Xiao, 2013; Mantovani, Leckman, Grantz, King, Sporn, Lisanby, 2007). Consequently, there is considerable interest currently in the use of NIBS techniques to develop safe and effective new treatments for a range of neurological and psychiatric conditions, including movement disorders such as PD and TS. However, these techniques are often not suitable for use outside of the laboratory or clinic, or for use with young children. For this reason we chose to investigate the potential for the therapeutic use of *peripheral* nerve stimulation. Specifically, we investigated whether we could use rhythmic median nerve stimulation (MNS) to entrain those brain oscillations that have been linked to the initiation and suppression of movement, specifically the Beta rhythm (Engel & Fries, 2010; Schnitzler & Gross, 2005). This work was motivated by two recent reports. First, the direct prediction arising from a recent computational model that the occurrence of motor tics in TS could be effectively reduced by externally stimulating the sensorimotor cortex (Caligiore, Mannella, Arbib, Baldassarre, 2017) and second, the recent empirical finding that compensatory alterations in TS may be associated with increased beta synchronization in patients with TS. Specifically, the finding that increased beta power was associated with reduced tic severity in TS (Niccolai, van Dijk, Franzkowiak, Finis, Südmeyer, Jonas, et al., 2016).

Oscillations of the brain’s electromagnetic activity are thought to reflect the synchronised firing of populations of neurons and it is known that GABA-mediated interneurons play a key role in co-ordinating the synchronised activity of populations of pyramidal neurons that give rise to brain oscillations (Schnitzler & Gross, 2005). Beta oscillations have long been associated with sensorimotor function (Armstrong, Sale, Cunnington, 2018), are linked to maintaining the current motor set (Engel & Fries, 2010), and become de-synchronised when a movement is initiated (Armstrong, Sale, Cunnington, 2018). As noted above, the beta (15-30Hz) frequency band is particularly relevant to the occurrence of tics in TS. Also, it is noteworthy that studies of EEG signals that are thought to arise in the supplementary motor area (SMA) ahead of movement indicate that these signals are abnormal in individuals with TS ahead of tic execution (Obeso, Rothwell, Marsden, 1981).

To investigate whether peripheral MNS could be used to entrain movement-related brain oscillations, we adapted an approach used by Thut et al. (2011) in which they had reported that rhythmic pulses of transcranial magnetic stimulation [TMS] could be used to entrain cortical alpha (8-14Hz) oscillations. In their study, Thut and colleagues delivered five-pulse rhythmic trains of TMS at each individual’s preferred alpha frequency (a-TMS) and demonstrated alpha entrainment (i.e., increased alpha power and phase synchrony). Importantly, this effect was not observed in a control condition in which five-pulse trains of TMS were delivered arrhythmically within the same time period.

In our study we combined electroencephalographic (EEG) recording with 10-pulse trains of rhythmic beta-band (19Hz) MNS versus 10-pulse trains of arrhythmic MNS. We demonstrate that *rhythmic* but not *arrhythmic* trains of MNS lead to entrainment of beta oscillations (i.e., sustained increase in 18-20Hz power and phase synchrony) contralateral to the site of peripheral stimulation, and that rhythmic and arrhythmic MNS lead to statistically different after effects once stimulation has ceased.

## Methods

### Participants

Twenty healthy young adults (8 males) aged between 20 and 35 years (mean age of 25.2 years) took part in the study. All of the subjects were right-handed. There were no requirements to take part in the study. Participants were given 8 pounds for their time. None of the recruited participants withdrew from the study. All participants signed the consent form of the study, approved by the Ethics Committee in Nottingham University.

### Median nerve stimulation

The right median nerve was stimulated using Digitimer DS7A HV Current Stimulator (*Digitimer Ltd, Hertfordshire, Uk*). Stimulation was delivered using a bar electrode consisting of two stainless steel disk electrodes with a diameter of 8mm separated by 30mm placed on the right wrist of the participants. The intensity of the stimulation used was the minimum intensity at which a thumb twitch was seen. The stimulator was controlled by a bespoke MATLAB vR2017a script on a Mac Pro computer running High Sierra (v. 10.13.6).

### EEG recording

EEG data was recorded from 64 electrodes using a BioSemi Active Two system. EEG data was recorded with a sampling rate of 1024Hz, this was later down-sampled to 128Hz. The impedance of the electrodes was kept under 30μV for all participants. Reference electrodes were placed on the left and right mastoids. Bipolar vertical and horizontal EOG electrodes were also recorded.

### Data Analysis

EEGlab (14.1.1) was used to pre-process and analyse the data. Data were low-pass filtered at 45Hz and high-pass filtered at 1Hz. Channels showing aberrant behaviour were deleted and noisy channels were interpolated. Automatic Artifact Removal (AAR) was used to remove EOG artifacts, using recursive least squares regression. Time-windows of -1 to 3 seconds, time-locked to the first pulse of the train were extracted. The whole second before the start of the stimulation was used as baseline. Epochs showing abnormal trends or excessive noise were rejected. Specifically, epochs showing a signal amplitude at +-100μV in one or more channels would be rejected; those epochs with signal slopes exceeding a threshold of 50μV in one or more channels would be rejected; those epochs with 5 times the standard deviation in the probability distribution would be rejected; those epochs which their kurtosis statistic was larger than 5 times of the standard deviation of the data would be rejected. Artifacts were detected by running Independent Component Analysis (ICA) and components were rejected with the use of Multiple Artifact Rejection Algorithm (MARA) and visual inspection. In total, based upon these criteria, only one channel was deleted from one participant. The average number of epochs between participants after epoch rejection was 129 in the rhythmic condition and 126 in the arrhythmic condition.

### Experimental procedure

300 trains, each consisting of 10 pulses of MNS, were delivered once every 4 seconds to the right median nerve of the participants while EEG was being recorded. The duration of each individual pulse was 200μs and the total duration of the pulse trains was 452ms. Pulse trains were delivered randomly in two different conditions as follows. In the rhythmic Beta-band MNS condition, pulses were delivered every 52ms (i.e. at a frequency of 19Hz); whereas in the arrhythmic condition, pulses were delivered during the same time window as the rhythmic condition (i.e., 452ms), but the intervals between individual pulses was not uniform and were selected pseudo-randomly for each pulse train, as illustrated in Figure 1.

**Figure 1.**
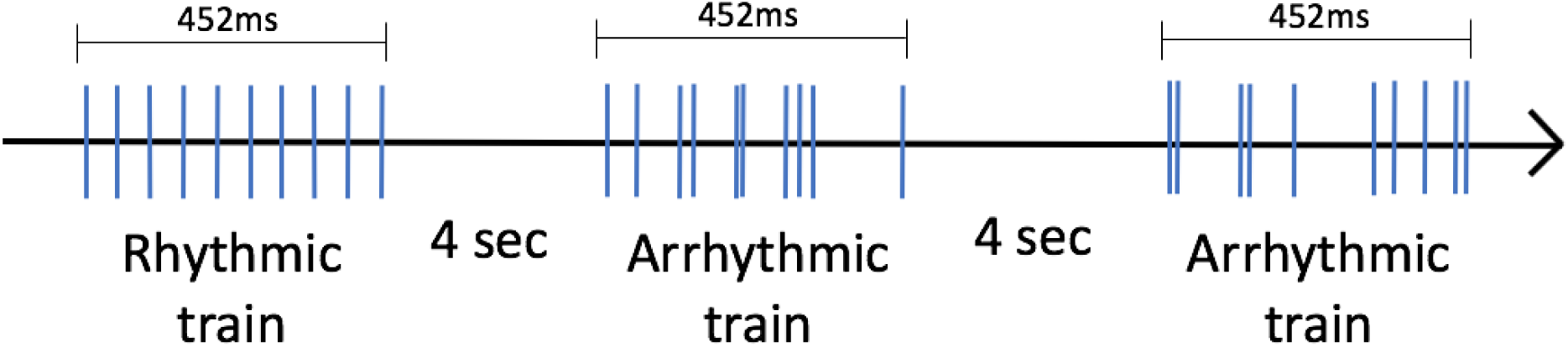
Illustrates the design of the study and difference between Rhythmic and Arrhythmic pulse trains.

Three short breaks were provided throughout the testing, one every 75 trains. Participants were instructed to stay still, relax, and keep their eyes open throughout. Participants were monitored throughout with a video camera to ensure that the thumb twitch was always present and the intensity of stimulation was adjusted if necessary. The starting intensity threshold was determined for each individual using a trial and error procedure of single pulses (ascending and descending) until a minimum threshold was established at which noticeable thumb twitch was reliably observed.

### Data Analysis

Analyses of the EEG data was performed to obtain event-related spectral perturbation (ERSP) values and phase-locking values (PLV) for a frequency range corresponding to 18-20Hz. Beta oscillations (13-30Hz) were plotted for every stimulation pulse and scalp maps were constructed for the entire scalp. Statistical analyses were performed using a one-tailed, paired-sample, t-test correcting for multiple comparisons using False Discovery Rate (FDR; Benjamini & Hochberg, 1995).

## Results

All analyses were performed on for the central scalp electrode (C1) located over the left sensorimotor cortical area.

### Event Related Spectral Perturbation (ERSP)

Time-frequency analysis indicated that there was an increase in ERSP at the beginning of the pulse train for both the rhythmic (19Hz) MNS and arrhythmic conditions that involved both Mu (8-12Hz) and beta (13-30Hz) bands (Figure 2). By contrast, after that initial increase in ERSP there was a sustained increase in the rhythmic MNS condition (Figure 2B) in a narrow band that peaked at the frequency of the stimulation (i.e. 19Hz). This increase in power centred at 19Hz was absent in the arrhythmic condition (see Figure 2A). This difference becomes clearer when subtracting the arrhythmic condition from the rhythmic condition (Figure 2C).

**Figure 2:**
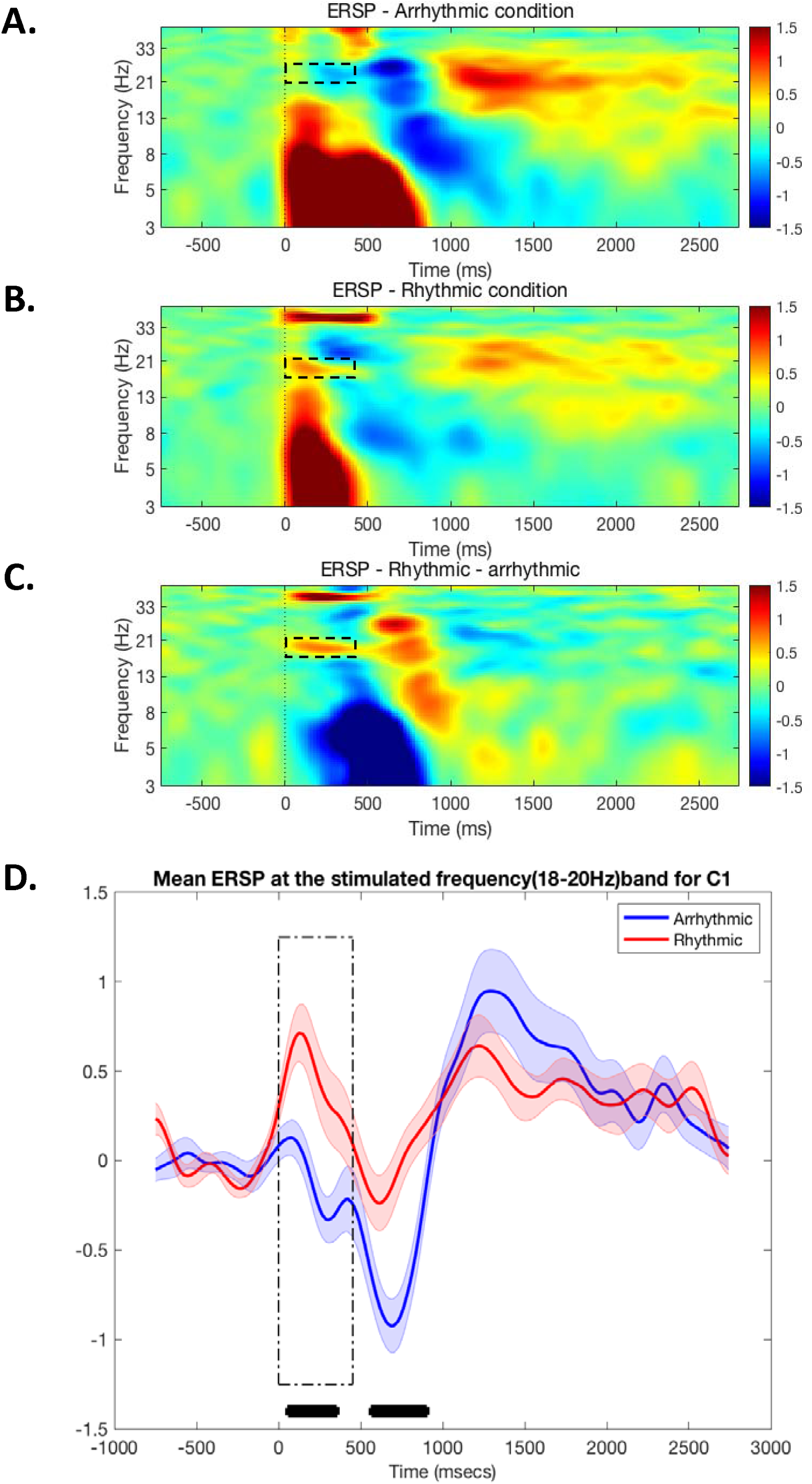
Time-frequency analyses of event-related spectral perturbation (ERSP) values at electrode C1 for **A**. Arrhythmic MNS condition. **B**. the Rhythmic 19Hz MNS condition. **C**. the subtraction of the Rhythmic – Arrhythmic conditions. In each case 0 on the x-axis indicates the onset of the pulse train. The broken rectangle indicates the 18-20Hz frequency range for the duration of the pulse train. **D**. A time-frequency plot of the average ERSP values within the frequency range 18-20Hz for the Rhythmic (red) and Arrhythmic (blue) MNS conditions. The broken rectangle indicates the duration of the stimulation and the black * symbols indicate individual time points at which the ERSP values were significantly different between conditions (p < .05 ^FDR-corrected^).

Figure 2D illustrates the average of ERSP values recorded from electrode C1 at 18-20Hz for the Rhythmic and Arrhythmic conditions. Inspection of this figure illustrates two observations of note. First, there is a significant increase in beta power at the frequency of the rhythmic stimulation relative to the arrhythmic condition [p < .05^FDR-corrected^] which occurs during the period of stimulation (from 78ms to 375ms). Second, there was a later, statistically significant (p < .05^FDR-corrected^), reduction in the average ERSP (desynchronization) for Arrhythmic condition compared to the Rhythmic condition which occurred after the stimulation had ceased. This result indicates that rhythmic beta-band stimulation appears to significantly reduce the magnitude of Beta-band desynchronization that is frequently been associated with movement initiation.

### Inter-trial coherence (ITC)

Analysis of inter-trial coherence (also referred to as phase-locking values) indicated that there was an increase in phase locking at the beginning of the pulse trains for both the Rhythmic and Arrhythmic MNS conditions that involved both Mu and Beta band frequencies (see Figure 3). Following this initial phase, the Rhythmic MNS condition exhibited an increase in ITC within a narrow band that peaked at the frequency of the stimulation (i.e. 19Hz) (Figure 3B). Importantly, this increase in phase locking centred at 19Hz was absent in the arrhythmic condition (Figure 3A). This difference becomes clearer when subtracting the Arrhythmic from the Rhythmic condition (see Figure 3C).

**Figure 3:**
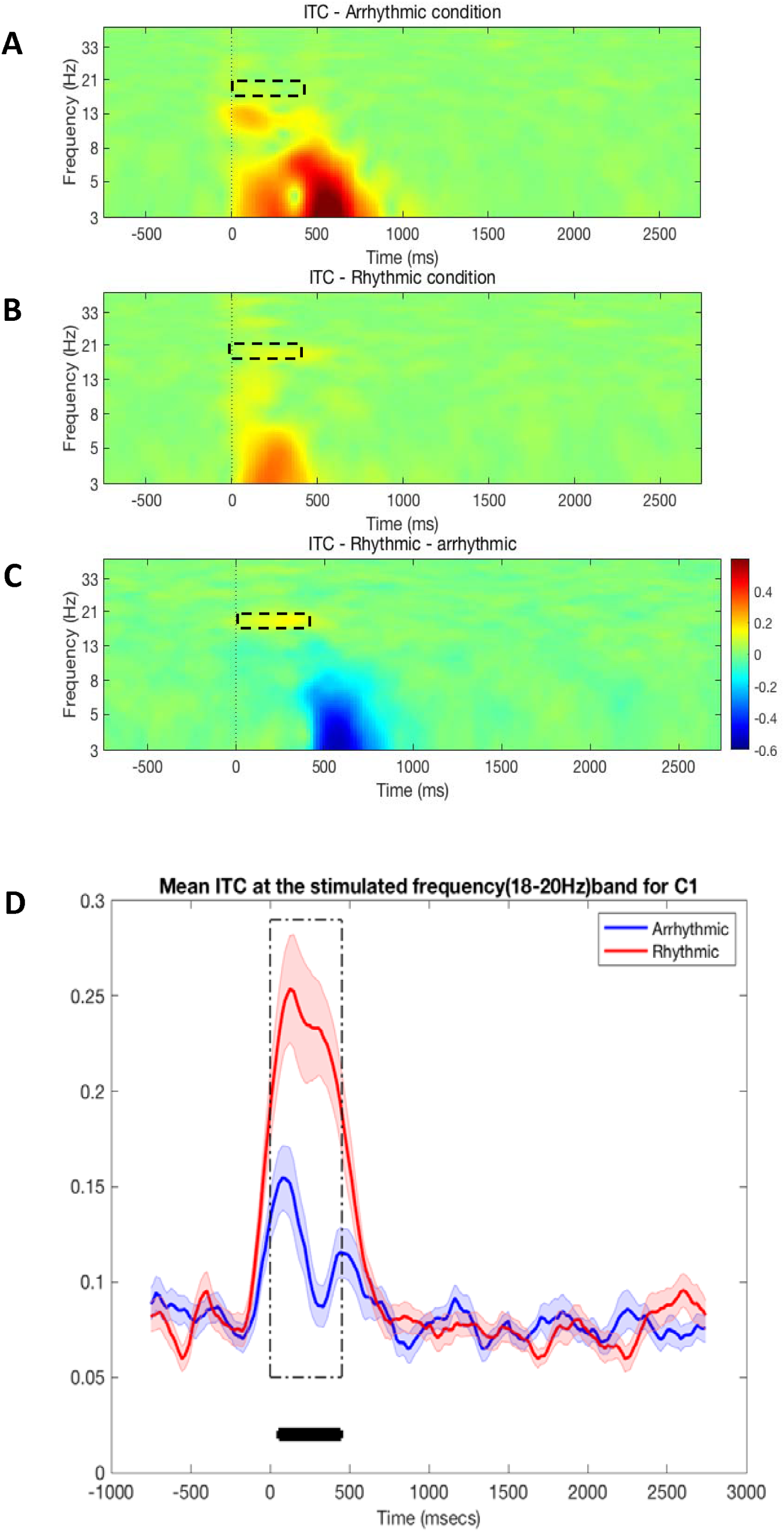
Time-frequency analyses of inter-trial coherence (ITC) values at electrode C1 for **A**. Arrhythmic MNS condition. **B**. the Rhythmic 19Hz MNS condition. **C**. the subtraction of the Rhythmic – Arrhythmic conditions. In each case 0 on the x-axis indicates the onset of the pulse train. The broken rectangle indicates the 18-20Hz frequency range for the duration of the pulse train. **D**. A time-frequency plot of the average ITC values within the frequency range 18-20Hz for the Rhythmic (red) and Arrhythmic (blue) MNS conditions. The broken rectangle indicates the duration of the stimulation and the black * symbols indicate individual time points at which the ITC values were significantly different between conditions (p < .05 ^FDR-corrected^).

Figure 3D shows the mean ITC values for the 18-20Hz frequency range recorded at electrode C1 for each conditions. Inspection of this figure reveals that there is a sustained increase in ITC that is specific to the time period of the MNS stimulation (from 54ms to 460ms) and is only observed for the Rhythmic 19Hz MNS condition (p < .05^FDR-corrected^). Importantly, and in contrast to the ERSP data reported above, there is no significant difference between the MNS conditions that outlasts the period of stimulation. This suggests that the increased in frequency-specific coherence is very likely driven by the individual pulses of the rhythmic pulse train. This will be explored below.

### Hemispheric specificity of rhythmic MNS entrainment

To examine any hemispheric specificity for Beta entrainment effects, we examined mean ERSP values within the stimulated frequency range (18-210Hz) recorded at the left hemisphere scalp contralateral to the peripheral median nerve stimulation (electrode C1), and the right hemisphere scalp ipsilateral to the stimulation (electrode C2). The results of this analysis is shown in Figure 4. Inspection of this figure illustrates that there is an initial increase in ERSP values at the beginning of the Rhythmic MNS pulse train at both the contralateral and ipsilateral electrode sites. However, this is followed by a sustained increase in ERSP values at the contralateral electrode site (C1) compared to the ipsilateral electrode site (C2) [p < .05^FDR-corrected^). Importantly, this increase in power (ERSP) recorded over the contralateral scalp is sustained after the Rhythmic MNS stimulation has ceased (i.e., from 984ms to 1265ms, from 1476ms to 1687ms and from 2093ms to 2531ms: see Figure 4).

**Figure 4:**
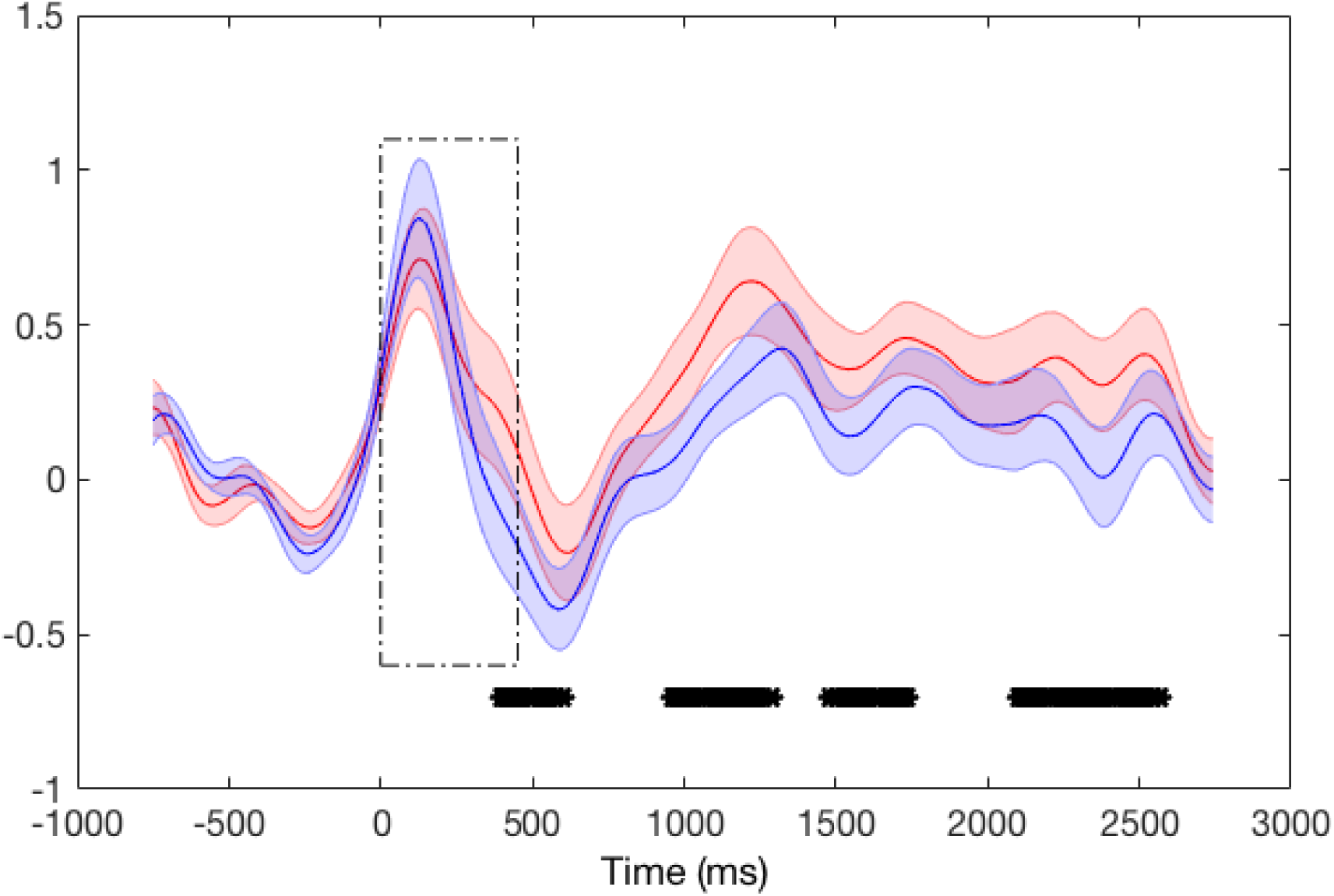
Illustrates a time-frequency plot of the average ERSP values within the frequency range 18-20Hz for the contralateral electrode site C1 (red) and the ipsilateral electrode site C2 (blue) for the Rhythmic 19Hz MNS condition. The broken rectangle indicates the duration of the stimulation and the black * symbols indicate individual time points at which the ERSP values were significantly different between conditions (p < .05 ^FDR-corrected^).

### Phase re-setting in response to each MNS pulse

A previous study had demonstrated that rhythmic TMS pulses delivered at Alpha-band frequencies resulted in a progressive synchronization of the targeted brain oscillation frequency and that enhanced synchronization was critically dependent on the existing, pre-TMS, phase of the signal generator (Thut et al., 2011). Importantly, in this study it was demonstrated that each individual TMS pulse had the effect of re-setting that phase. To determine if Rhythmic MNS has a similar effect of re-setting the phase of the targeted signal generator, we plotted the mean Beta-band (13-30Hz) oscillations for each pulse within the MNS pulse train for both rhythmic and arrhythmic stimulation (Figure 5A). The results indicate that rhythmic MNS leads to the predicted re-setting of the phase for every pulse of the train. By contrast, this phase-reset is only seen for the first 3 pulses in the arrhythmic condition. This is consistent with our observation of an increase in power and ITC that is observed at the beginning of the pulse train for arrhythmic MNS (pulses 1-3), and with the finding that there is a sustained increase in power and ITC, centred at 19Hz, for pulses 4-10 only during rhythmic MNS.

**Figure 5:**
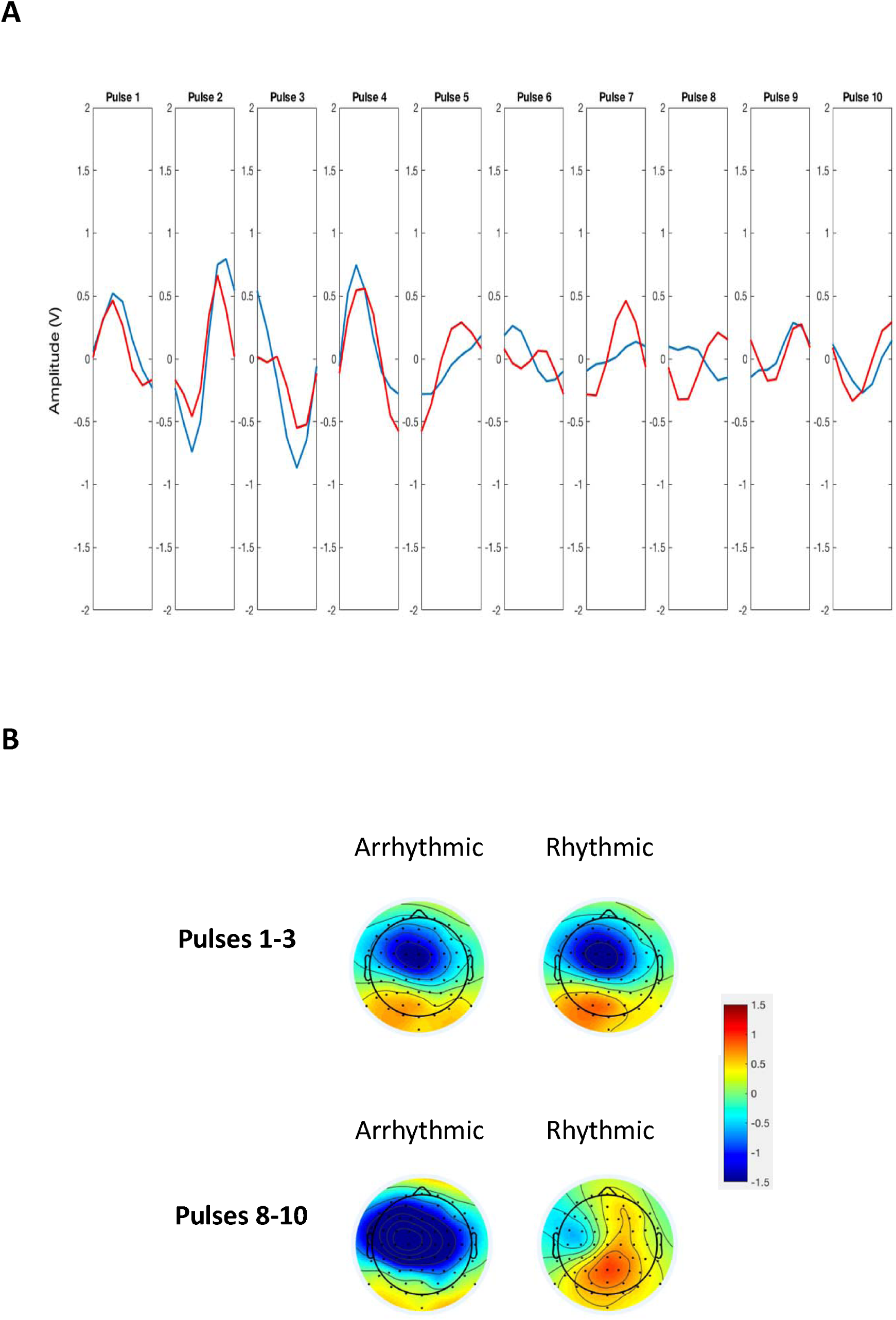
**A**. Illustrates the mean Beta-band (13-30Hz) oscillations following each of the ten pulses within the pulse train for both Rhythmic (red) and Arrhythmic (blue) MNS. **B**.

### Topography

Scalp maps of the event-related potentials (ERPs) for Beta-band (13-30Hz) were plotted using all electrodes, separately for pulses 1-3 (0-150ms) and pulses 8-10 (350-500ms), for both Rhythmic and Arrhythmic conditions. The results are presented in Figure 5B. Analysis of these scalp topographies revealed that for pulses 1-3 there was no significant difference between conditions at any of the 64 electrode positions (Figure 5B: upper panel). By contrast, for pulses 8-10 the scalp topographies are quite different (Figure 5B: lower panel) and the maps for each condition differ significantly (p<0.05) from one another at 57 (from 64) electrode locations (only electrodes P7, P9, P07, O1, Iz, P10, TP7 were not significantly different).

### After-effects of the stimulation

To examine the after effects of stimulation, including after effects observed at frequencies outside of the targeted stimulation frequency, we examined average high Beta (21-30Hz) power (ERSP) and Mu (8-13Hz) band power following Rhythmic (19Hz) MNS compared to Arrhythmic MNS. Figure 6A shows time-frequency plots for the Rhythmic (left panel) and Arrhythmic (right panel) conditions. Inspection of this figure indicates that there is a larger Beta and Mu desynchronization (∼1000ms) effect, and Beta-rebound effect (increased synchronization) (∼1250ms) effect, for the Arrhythmic MNS condition compared to the Rhythmic (19Hz) MNS condition. This is important as it suggests that the event-related Mu and Beta desynchronization, and Beta-rebound, effects that are associated with movement initiation, can be significantly reduced using rhythmic Beta-band entrainment. Figure 6B and 6C show the time course of this effect for Beta and Mu power respectively, and indicates time points where the difference between conditions is statistically significant (p < .05^FDR-corrected^).

**Figure 6:**
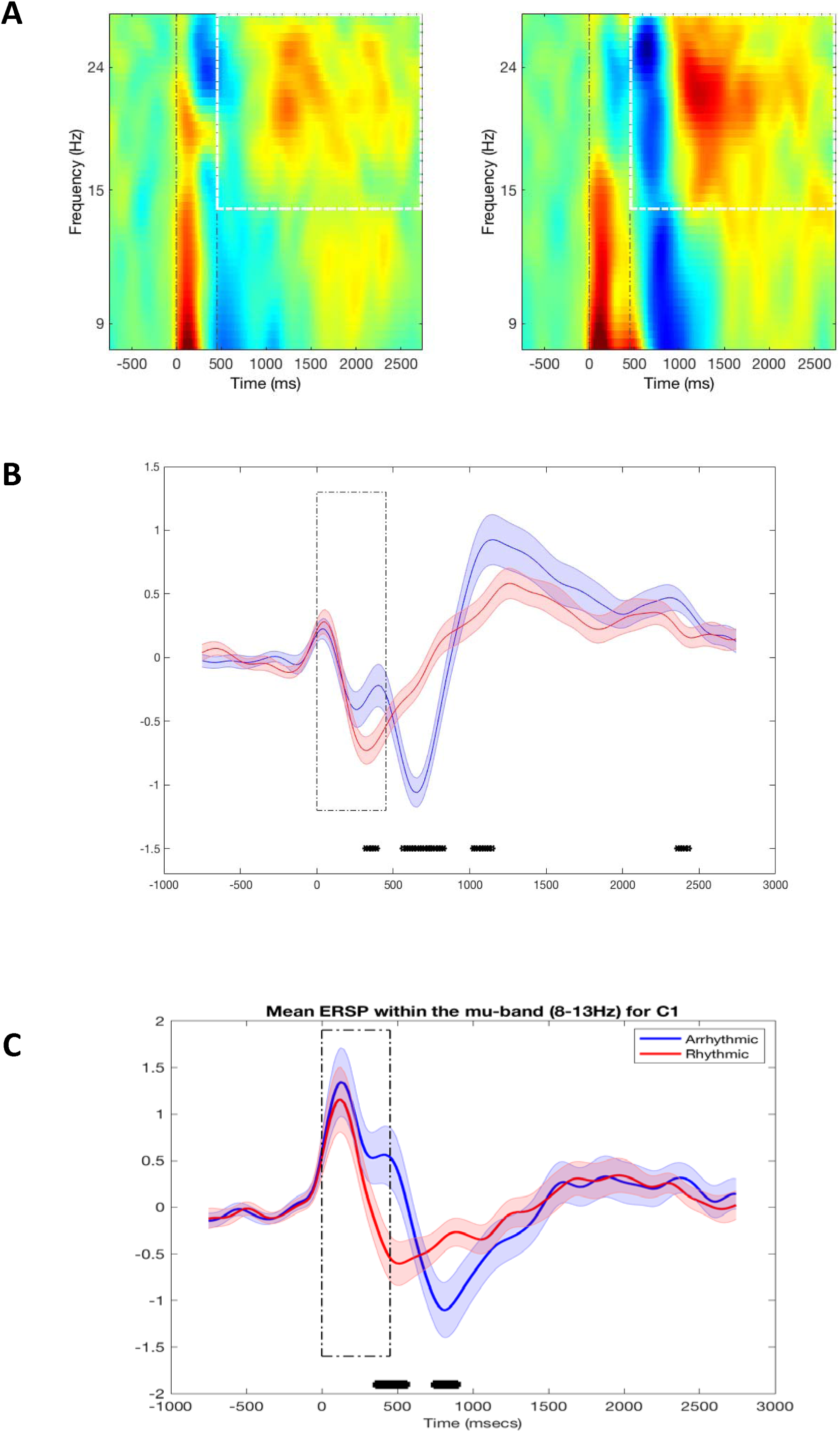
**A**: Illustrates a time-frequency plot of the average ERSP values within the frequency range 18-20Hz measured at electrode C1 for the Rhythmic (19Hz) MNS condition (left panel) and the Arrhythmic MNS condition (right panel). 0 on the x-axis indicates the start of the MNS pulse train and the broken white dotted vertical line (at 452ms) indicates the end of the pulse train. The broken white rectangle indicates the Beta-band frequency range after the period of MNS has ceased. **B**. Average ERSP values within the Beta frequency range (21-30Hz) for the Rhythmic (19Hz) MNS (red) and Arrhythmic MNS (blue) conditions. The broken black rectangle indicates the duration of the MNS pulse train. The black * symbols indicate individual time points at which the ERSP values were significantly different between conditions (p < .05 ^FDR-corrected^). **C**. Average ERSP values within the Mu frequency range (8-13Hz) for the Rhythmic (19Hz) MNS (red) and Arrhythmic MNS (blue) conditions. The broken black rectangle indicates the duration of the MNS pulse train. The black * symbols indicate individual time points at which the ERSP values were significantly different between conditions (p < .05 ^FDR-corrected^).

## Discussion

We investigated whether peripheral nerve stimulation - specifically rhythmic stimulation of the median nerve at the right wrist - could be used to entrain brain oscillations linked to the initiation and suppression of movements. We delivered rhythmic 10-pulse trains of MNS at a frequency of 19Hz while recording EEG, and compared this against a control condition in which we delivered 10-pulse MNS trains arrhythmically within the same time window. The results of this study can we summarised as follows. First, we demonstrate that rhythmic 19Hz MNS leads to a sustained, statistically significant, increase in beta (18-20Hz) power and phase-synchrony (ITC) over the sensorimotor cortex contralateral to the site of MNS delivery. One important difference between the power (ERSP) and phase-synchrony (ITC) measurements is that significant increases in ITC were only observed during stimulation whereas significant increases in power following rhythmic MNS outlasted the period of stimulation. Second, we demonstrate that, consistent with previous reports of the effects of rhythmic TMS (Thut et al., 2011), the effects of rhythmic MNS can be divided into two distinct phases. An initial period (pulses 1-3), in which there is a broadband increase in power and phase-synchrony that is similar for both rhythmic and arrhythmic MNS, and is observed contralaterally and ipsilaterally to the site of peripheral stimulation. This initial period is followed by a later period (pulses 4-10) in which there is a sustained increase in power and phase-synchrony for the rhythmic MNS condition that is largely observed over the contralateral sensorimotor cortex. Third, we demonstrate that increased phase-synchrony observed following rhythmic MNS was likely due to sustained effects of phase re-setting in response to each MNS pulse within the 10-pulse train. Fourth, we demonstrate that the during the initial phase (pulses 1-3) of MNS the spatial topography for the rhythmic and arrhythmic conditions did not differ. By contrast, during the later period (pulses 4-10) there were large, statistically significant differences in scalp topography. Finally, and of some importance, we demonstrate that rhythmic MNS leads to sustained after-effects that outlast the period of stimulation. Specifically, rhythmic 19Hz MNS leads to a significant *reduction* in the beta-band desyncronization that is most often associated with stimulation of the sensorimotor cortex (Jasper and Penfield, 1949), including the initiation of movement (Engle & Fries, 2010), and a *reduction* in the magnitude of the subsequent beta-rebound effect (Neuper and Pfurtscheller, 2001) which has been proposed as an integrative signal that exerts an inhibitory influence over the primary sensorimotor cortex (Donner and Siegel, 2011). These results are discussed below.

### Alterations in beta power and phase synchrony following rhythmic 19Hz MNS

As noted above, Thut et al. (2011) had demonstrated that rhythmic 5-pulse trains of TMS delivered at alpha-band frequencies led to an increase in power and phase-synchrony at the stimulated alpha frequency that was not observed following 5-pulse TMS trains of arrhythmic stimulation. This effect was characterized by an initial broad band increase in EEG power that involved both mu and beta frequency bands and was seen for both rhythmic and arrhythmic stimulation, which was then followed by a sustained increase in EEG power and phase-synchrony that was focused at the targeted frequency of alpha stimulation and only observed for rhythmic stimulation. The results of the current study indicate very clearly that highly similar effects can be obtained using 10-pulse trains of rhythmic MNS delivered at 19Hz.du

An important, finding from the Thut et al. (2011) study was that the increase in phase-synchrony observed in their study during rhythmic TMS was due to the phase re-setting effect of each individual TMS pulse, which led to a sustained and progressive synchronization of the targeted brain oscillation frequency for rhythmic TMS pulse trains but this was not sustained for arrhythmic TMS pulse trains. This indicates that while each TMS pulse is capable of re-setting the phase of the underlying oscillatory generator, it is only when these TMS pulses occur at a specific frequency that you observe a sustained increase in phase-synchrony for the duration of the pulse train.

The demonstration that brain oscillations associated with movement initiation/suppression may be entrained by using external stimulation delivered at a target frequency has a number of important implications for influencing motor behaviour, including motor behaviour linked to brain health conditions associated with unwanted movements such as TS. Consistent with this view, previous studies have demonstrated that: volitional movements executed during periods of increased beta-band power are significantly slowed (Gilbertson, Lalo, Doyle, Di Lazzaro, Cioni, Brown, 2005); beta entrainment using 20Hz transcranial alternating current stimulation (tACS) can significantly impact upon the kinematics of naturalistic movements (Pogosyan, Gaynor, Eusebio, Brown, 2009); and, that tACS delivered at the frequency of the resting tremor of PD patients, but out of phase with the tremor, can reduce resting tremor amplitude by an average of 50% (Brittain, Probert-Smith, Aziz, Brown, 2013).

### Aftereffects of rhythmic MNS

One un-anticipated finding arising from this study was that, in addition to inducing increases in power and phase-coherence at the targeted beta frequency, rhythmic MNS also induced significantly different after effects compared to the arrhythmic MNS control condition, including after effects at brain oscillation frequencies that had not been directly stimulated. Specifically, following rhythmic 19Hz stimulation there was a significantly reduced reduction in average power (desynchronization) in both mu (8-13Hz) and high-beta (21-30Hz) frequency bands, that was followed by a significant reduction in the subsequent increase in high-beta power rebound (synchronization) effect.

It should be noted that motor-related beta oscillations are thought to reflect the maintenance of the current motor set (i.e., a stable motor state) and the suppression of states associated with new movements. By contrast, movement preparation and execution is associated with the attenuation of beta oscillations (Engle & Fries, 2010). Furthermore, alterations in beta oscillations have been reported in brain health conditions linked to movement disorder, with *increased* beta-band oscillations linked to akinesia in PD (Brown, 2007) and decreased beta-band oscillations associated with motor tics in TS (Israelashvili, Loewenstern, Bar-Gad, 2015). For this reason, our finding that bursts of rhythmic peripheral nerve stimulation may be sufficient to induce periods of altered beta-band power that outlast the period of direct stimulation is of considerable interest, particularly if it can be demonstrated that such effects may be additive leading to a sustained period of increased beta-band power.

## Acknowledgements

This work was supported by research grants from Tourettes Action and by the NIHR Nottingham Biomedical Research Centre. The views expressed are those of the authors and not necessarily those of the NHS, the NIHR or the Department of Health. We thank Tourettes Action (UK) for assisting with participant recruitment.

